# Uncertainties in tumor allele frequencies limit power to infer evolutionary pressures

**DOI:** 10.1101/134049

**Authors:** Javad Noorbakhsh, Jeffrey H. Chuang

## Abstract

We read with great interest the paper by Williams, *et. al.*[1], who argued for neutral evolution in tumors by analyzing The Cancer Genome Atlas (TCGA) data. They supported this conclusion by showing high *R*^2^ values for fits to a neutral evolution model predicting *M* ∝ 1 / *f*, where *M* is the number of somatic mutations with allele frequency ≥ *f*. However, we believe a conclusion of neutrality must be treated cautiously, as high *R*^2^ values are consistent with many evolutionary models.

---

For example, we analyzed phenomenological models similar to that of [1] but with parameter *k*, such that *M* ∝ 1/ *f* ^*k*^. Here *k* = 1 corresponds to the neutral model, *k* > 1 corresponds to diversifying selection (excess of rare mutations), and *k* < 1 corresponds to purifying selection (excess of high frequency mutations). We reanalyzed the TCGA data to determine whether values other than *k* = 1 fit the data better. To reduce pipeline uncertainties we used only tumors for which calls were made by Mutect [2], and similar to [1] we only used mutations with read count *≥*10, and alternative read count *≥*3, and only analyzed tumors with *≥*5 genes within the fitting range (0.12 *< f <* 0.24). We then reproduced Figure 3 from [1] by fitting mutation count to 1/ *f* (Figure 1). Our values of *R*^2^ were high though not identical to [1], which is due to differences in tumor sets and the lack of information about exact procedures used in [1]. To determine whether the fit was due to neutral evolution, we repeated the same analysis by fitting to the functions 1/ *f* ^2^ (diversifying selection) and 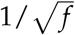 (purifying selection) (Figure 1). In all cases, we were able to closely fit the TCGA data (mean *R*^2^ values were 0.84, 0.87, 0.74 for *k* = 1, 0.5, 2), but the purifying selection model 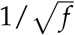 in fact fit the data slightly better. Although our analysis does not clearly show a lack of neutrality, it does indicate that *R*^2^ is not a good measure for distinguishing neutral evolution.

**Figure 1:**
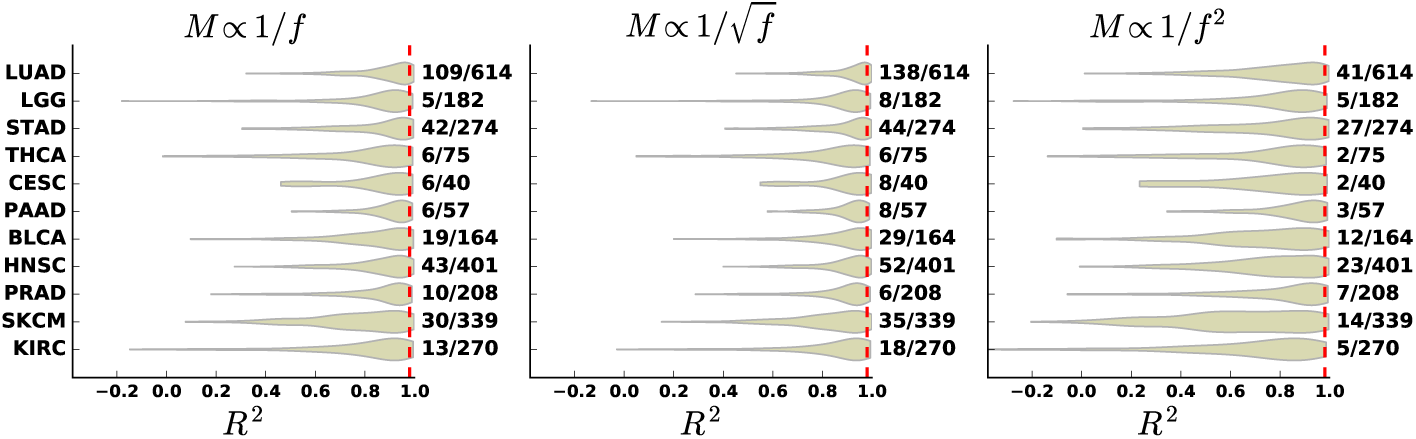
Distribution of *R*^2^ for fits of TCGA allele frequency distribution data to three different models. The numbers on the right side of each plot show the fraction of total tumors in each cancer type with *R*^2^ *>* 0.98 (right side of red dashed line).

A more fundamental consideration is that noise inherent in the *M*(*f*) curves limits conclusions about neutrality. Assuming the true allele frequency of a mutation is *f*_*True*_, the observed allele frequency *f*_*Obs*_ will be a sample from a binomial distribution with mean *μ* _*f*_ = *f*_*True*_ and standard deviation *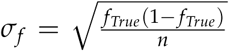*, given read depth *n* (on average *n* = 102 in the TCGA samples). In the fitting range 0.12 *< f*_*True*_ *<* 0.24, *σ*_*f*_ can take on values as large as 0.04, i.e. *∼* 30% of the fitting range. We analyzed the effect of this noise directly by simulating observed *M*(*f*) curves according to underlying neutral (*k* = 1), purifying (*k* = 1/2), and diversifying (*k* = 2) selection models. *M*(*f*) curves were generated by sampling values of *f*_*True*_ from the underlying model and then for each value reporting an *f*_*Obs*_ generated from the binomial distribution with mean *f*_*True*_ and read depth *n*, where *n* was drawn from a lognormal fit to the pooled TCGA read depth distribution. Figure 2 shows randomly generated *M* curves obtained by resimulating this process, suggesting that measurement uncertainty can signficantly impact the shape of the observed curve and obscure the underlying evolutionary process. Moreover, we repeatedly simulated *M*(*f*) curves for each generating process (*k* = 1/2, 1, 2) and tested whether the true generating process could be identified. Mean and standard deviation of *R*^2^ values are shown in Table 1. *R*^2^ values to the true model (diagonal elements) are only marginally better than to the incorrect models, and in all cases these differences are less than the standard deviation across replicates, suggesting that *R*^2^ is not a sensitive measure for resolving the evolutionary process.

**Figure 2:**
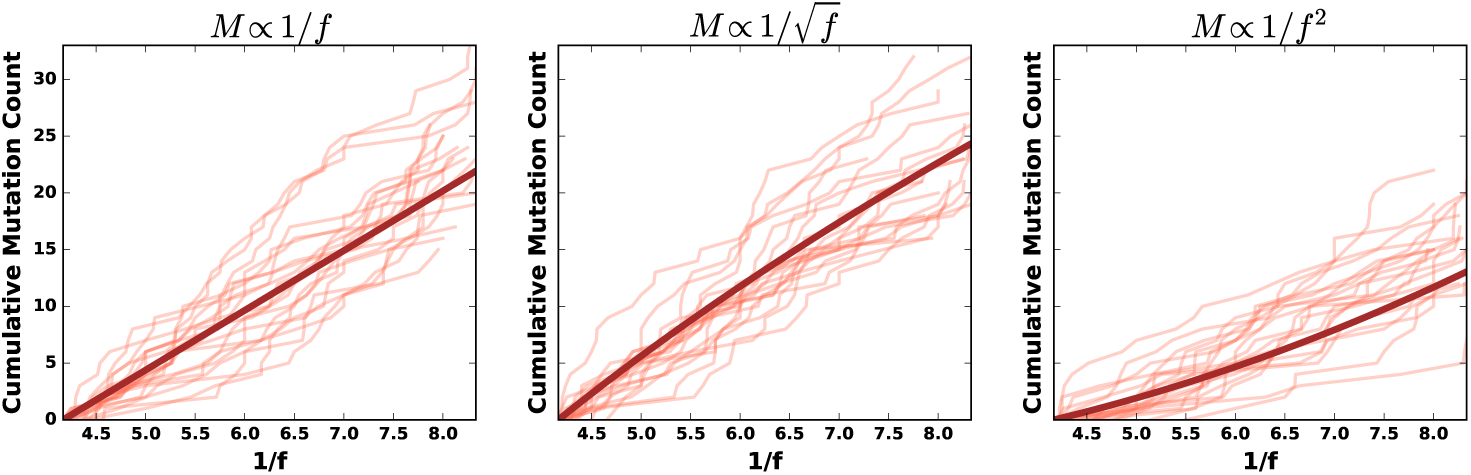
Simulated allele frequency distributions for different generating processes. Thin curves are individual examples of simulated *M* curves from the (left) neutral, (center) purifying selection, and (right) diversifying selection processes, while the thick curves are the ideal when no measurement noise exists.

**Table 1.** Fits of simulated data from neutral (1/ *f*), purifying selection 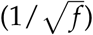, and diversifying selection (1/ *f* ^2^) to the expected *M* curves for all three processes. Mean *R*^2^ values are shown, with standard deviations shown in parentheses.

| Fit to : | | $1/f$ | $1/\sqrt{f}$ | $1/f^2$ |
| --- | --- | --- | --- | --- |
| Generating Process : | $M \propto 1/f$ | 0.95 (0.04) | 0.95 (0.04) | 0.93 (0.06) |
| | $M \propto 1/\sqrt{f}$ | 0.94 (0.05) | 0.95 (0.04) | 0.90 (0.09) |
| | $M \propto 1/f^2$ | 0.94 (0.05) | 0.93 (0.05) | 0.94 (0.05) |

Williams, *et. al.* have provided a valuable conceptualization of population dynamics in tumors and shown that neutrality is possible. However, models with selection can provide similarly good fits to the TCGA data, and TCGA data still yield substantial uncertainties about the true frequency distributions. More refined evolutionary models and further increases in sequencing depth will be important for resolving the balance of selection and neutrality in cancer.

Research reported in this publication was supported by the National Cancer Institute under award numbers P30CA034196 and R21CA191848. The content is solely the responsibility of the authors and does not necessarily represent the official views of the NIH. JHC also acknowledges support from the Hope Foundation.

## References

1. Marc J Williams, Benjamin Werner, Chris P Barnes, Trevor A Graham, and Andrea Sottoriva. Nature genetics, 48(3):238–244, 2016.

2. Kristian Cibulskis, Michael S Lawrence, Scott L Carter, Andrey Sivachenko, David Jaffe, Carrie Sougnez, Stacey Gabriel, Matthew Meyerson, Eric S Lander, and Gad Getz. Nature biotechnology, 31(3):213–219, 2013.

